# Cardiomyocyte-intrinsic SLC25A1 regulates cardiac differentiation and mitochondrial function

**DOI:** 10.64898/2026.03.06.710200

**Authors:** Chiemela Ohanele, Nahum F. Arefeayne, Nasab Ghazal, Ethan H. Liu, Nina C. Turcanu, Matthew J. Horchar, Biplab Dasgupta, Adriana Harbuzariu, Chunhui Xu, Victor Faundez, Jennifer Q. Kwong

**Author notes:** Correspondence Emory University School of Medicine Department of Pediatrics 1750 Haygood Dr. NE Rm 432 Atlanta, GA 30322.

## Abstract

Cardiac morphogenesis is an intricate process that requires a precise coordination between metabolic and structural maturation, but how these processes are linked remain unclear. In previous work, we identified one candidate underlying this connection: the mitochondrial citrate carrier (SLC25A1), a critical regulator of embryonic heart development. Here, using systemic and cardiomyocyte-specific *Slc25a1* deletion in mice together with SLC25A1 knockout (KO) human induced pluripotent stem cell-derived cardiomyocytes (hiPSC-CMs), we demonstrate that SLC25A1 functions cell-autonomously within cardiomyocytes to regulate differentiation, mitochondrial maturation, and ventricular morphogenesis. Transcriptomic analysis of SLC25A1-deficient hearts revealed dysregulation of gene programs regulating cardiomyocyte differentiation and mitochondrial function. Consistent with these changes, loss of SLC25A1 in developing cardiomyocytes impaired mitochondrial function and resulted in defective ventricular wall compaction in vivo. Likewise, SLC25A1 KO hiPSC-CMs exhibited defective cardiomyocyte differentiation, disorganized myofibrils, and immature mitochondrial organization and function in vitro. Together, our findings position SLC25A1 as a cardiomyocyte-intrinsic, cell-autonomous regulator that links mitochondrial citrate export to developmental gene programs, revealing a mitochondrial regulatory axis for cardiomyocyte maturation and cardiac morphogenesis that contributes to congenital heart disease.

## Introduction

Cardiac morphogenesis is a complex process driven by both intrinsic pathways from within the developing myocardium as well as extrinsic regulation from the fetal environment. As the developing heart progresses through critical morphologic transitions such as looping, chamber formation and septation, coordinated transcriptional and metabolic reprogramming drive stage-specific growth and maturation ^1–4^. Disruptions to these processes can lead to congenital heart defects (CHD), which are the most common form of birth defect, occurring in approximately 1% of live births ^5^. To date however, only 20% of CHDs have a known cause ^6,7^. Therefore, defining the molecular mechanisms required for proper cardiac development is needed to connect genotype to phenotype and to inform future therapy development.

The mitochondrial citrate carrier (SLC25A1) is a mitochondrial inner membrane transporter responsible for citrate export to the cytosol ^8^. Because citrate is a precursor for cytosolic acetyl-CoA production, SLC25A1 thus links mitochondrial metabolism to pathways like lipid biosynthesis and epigenetic regulation ^2,9–13^. Our recent work demonstrated that SLC25A1 is required for cardiac morphogenesis and metabolic maturation in vivo ^14^. With systemic SLC25A1 loss, both hemizygous and homozygous deletion, produced CHDs, including ventricular septal defects, right ventricular hypoplasia, and ventricular noncompaction, with an increased incidence and severity of defects in full knockouts, indicating SLC25A1 haploinsufficiency for cardiac development. At the molecular level, SLC25A1 deletion disrupted cardiac mitochondrial ultrastructure and function, and altered epigenetic control of metabolic gene programs, consistent with a role for SLC25A1 in metabolic reprogramming during cardiac morphogenesis ^14^. Together, these data identify SLC25A1 as a key regulator of mitochondrial and developmental programs in the embryonic heart.

While our study identified structural cardiac defects resulting from systemic *Slc25a1* deletion, recent work has also implicated SLC25A1 in the regulation of the placenta–fetal heart axis ^15^. In that study, systemic *Slc25a1* knockout embryos displayed both placental malformations and CHDs, and selective deletion of *Slc25a1* in placental trophoblasts (cells important for nutrient storage and delivery), was sufficient to recapitulate the cardiac defects ^15–19^. These findings underscore the importance of the placenta in supporting fetal cardiac morphogenesis. However, although *Slc25a1* was deleted in cardiomyocytes and no overt postnatal cardiac abnormalities were observed, embryonic myocardial phenotypes were not examined in detail ^16^, leaving unresolved whether SLC25A1 is required for embryonic myocardial growth, differentiation, and maturation. Given its high expression in the developing myocardium and its established role in regulating mitochondrial metabolism and function across multiple cell types ^2,14,20,21^, we hypothesized that SLC25A1 functions cell-autonomously within cardiomyocytes to regulate morphogenesis and mitochondrial maturation, independent of placental effects.

Here, we tested the hypothesis that SLC25A1 exerts a cardiomyocyte-intrinsic role in regulating cardiac morphogenesis. Using both systemic and cardiomyocyte-specific knockout mouse models, we defined the requirement for SLC25A1 within the developing myocardium and distinguish cell-autonomous from placental contributions to CHD. In parallel, we employed human induced pluripotent stem cells (hiPSC) lacking SLC25A1 to examine its role in cardiomyocyte differentiation, mitochondrial organization, and respiratory function. Together, these studies identify SLC25A1 as a critical cardiomyocyte-intrinsic regulator of mitochondrial maturation and cardiomyocyte differentiation, providing mechanistic insight into an important axis of mitochondrial control of cardiac morphogenesis via SLC25A1 and the etiology of CHDs.

## METHODS

### Sex as a biological variable

Both male and female mice were included in this study. Although similar trends were observed in both sexes, the study was not powered to detect sex-specific differences, and analyses were therefore not stratified by sex.

### Animals

Slc25a1tm1a(EUCOMM)Wtsi knockout-first mice (*Slc25a1^+/-^*) were obtained from the Mutant Mouse Resource and Research Center at the University of California, Davis and maintained on a C57BL6/J background. *Slc25a^fl^*^/*fl*^ loxP-targeted mice were generated by crossing *Slc25a1^+/-^*mice to animals harboring a codon-optimized FLP recombinase (FLPo) under the control of the ubiquitously expressed phosphoglycerate kinase 1 (*Pgk1*) promoter [B6.Cg-Tg(Pgk1-flpo)10Sykr/J; The Jackson Laboratories, Strain #:011065] ^62^. *Slc25a1^fl/fl^;Tnnt2^Cre/+^* mice were generated by crossing *Slc25a1^fl^*^/*fl*^ mice to animals expressing a Cre recombinase under the control of the cardiomyocyte-specific rat cardiac troponin T promoter (The Jackson Laboratories, Strain #: 024240) ^63^.

Timed matings for the *Slc25a1^+/-^* line were conducted by intercrosses of 2–3-month-old *Slc25a1^+/-^* mice, with the morning of the detection of a copulation plug designated as E0.5. Timed matings for the *Slc25a1^fl/fl^;Tnnt2^Cre/+^* line were conducted by crossing 2–3-month-old *Slc25a^fl^*^/*fl*^ and *Slc25a1^fl/+^;Tnnt2^Cre/+^*mice, again with the morning of an observed copulation plug designated as E0.5. Genotyping of mice and embryos was performed on genomic DNA isolated from embryonic yolk sacs or tail biopsies.

### hiPSC culture and cardiomyocyte differentiation

hiPSCs were cultured under feeder-free conditions on Matrigel-coated tissue culture plates (Corning, 354277) in StemFlex medium (Gibco, A3349401) at 37 °C with 5% CO_2_ and passaged at 70-80% confluency using ReLeSR (StemCell Technologies, 100-0485). Cells were routinely tested and confirmed to be mycoplasma negative, assessed for pluripotency, and used for experiments at low passage.

hiPSCs were differentiated into cardiomyocytes by small molecule-guided Wnt modulation^64^. Undifferentiated hiPSCs were seeded onto Matrigel-coated 12-well plates and grown to approximately 95% confluency. Differentiation was initiated (day 0) by treatment with 6 μM CHIR99021 (Selleckchem, CT99021) in RPMI 1640 supplemented with B27 minus insulin. On day 2, medium was replaced with RPMI 1640 containing B27 minus insulin. On day 3, cells were treated with 5 μM IWR-1 (Sigma-Aldrich, I0161) in RPMI 1640 containing B27 minus insulin. From day 5 onward, cells were maintained in RPMI 1640 supplemented with B27 plus insulin, with medium changes every 2 days. Cells were harvested for downstream analyses between days 15 and 20 of differentiation, as indicated for individual assays.

### Generation and validation of SLC25A1 knockout hiPSC line

WT and SLC25A1 KO hiPSC lines were generated by the Emory Stem Cell and Organoids Core. SLC25A1 KO hiPSCs were derived from the parental female A18945 episomal hiPSC line (Thermo Fisher Scientific), which was originally generated from CD34⁺ human umbilical cord blood cells. Genome editing was performed using CRISPR/Cas9-mediated targeting of the SLC25A1 locus with a Gene Knockout Kit v2 (Synthego). The kit included three single-guide RNAs (sgRNAs) targeting SLC25A1: sgRNA1 (GGGAGCUAAGGCCGCGGUAC), sgRNA2 (CUCCCCCAGGGGACUGCGUG), and sgRNA3 (CCCCCUCCUCACCGAUGCCC).

Following genome editing, single-cell cloning was performed to isolate individual WT and SLC25A1-edited hiPSC clones. Candidate clones were screened by the Emory Integrated Genomics Core. Genomic DNA was amplified by PCR using Platinum SuperFi II PCR Master Mix (Invitrogen, 12368010) and primers flanking the sgRNA target sites (forward: GCATGTGGTTGCTGAGGAAC; reverse: TATGTTCCCCGCGGCAC). PCR products were subjected to Sanger sequencing at Genewiz (South Plainfield, NJ) using the sequencing primer GCATGTGGTTGCTGAGGAACTC. Sanger sequencing traces were analyzed using the Inference of CRISPR Edits (ICE) tool (Synthego) to determine editing efficiency and to identify clones harboring biallelic disruptive mutations at the SLC25A1 locus.

Two independent SLC25A1 KO clones (KO1 and KO2) and an isogenic WT control clone were selected and expanded for downstream analyses. Loss of SLC25A1 protein expression in KO clones was confirmed by immunoblotting.

Genomic integrity of edited and control hiPSC lines was assessed by G-banded karyotyping performed by WiCell. Pluripotency of WT and SLC25A1 KO hiPSC lines was verified by immunofluorescence staining for SSEA4, SOX2, TRA-1-81, and OCT4. Trilineage differentiation capacity was confirmed by immunostaining and flow cytometric analysis of ectodermal, mesodermal, and endodermal lineage-specific markers.

### Histology and immunostaining of embryonic heart and placenta

Placenta and hearts were fixed in 10% neutral-buffered formalin and embedded in paraffin. Paraffin-embedded tissues were sectioned (6 μm), deparaffinized, rehydrated, stained with hematoxylin and eosin (H&E), and imaged using an AxioScan 7 slide scanner (Zeiss). Placental layer thicknesses were measured on H&E-stained sections and normalized to whole placental thickness using ImageJ software (NIH) ^65^. Myocardial trabecular and compact layer thicknesses were measured on H&E-stained sections using QuPath image analysis software ^17^.

For immunofluorescence, paraffin-embedded sections were deparaffinized and rehydrated, and antigen retrieval was performed with Tris-EDTA buffer (pH 6.0). The following primary antibodies were used: anti-myosin heavy chain (Developmental Studies Hybridoma Bank, MF20, 1:20), anti-pHH3 (Cell Signaling Technologies, #9701, 1:100). Secondary antibodies used were: goat anti-mouse 488 (Southern Biotech, 103130, 1:200), and goat anti-rabbit 594 (Sigma, SAB4600107, 1:200). Slides were mounted with ProLong Gold Antifade Mountant with DAPI (Thermo Fisher Scientific, P36931) to counterstain nuclei. Fluorescence imaging was conducted using a Keyence BZ-X810 system or a Leica Stellaris confocal microscope.

### Immunocytochemistry of hiPSC-derived cardiomyocytes

hiPSC-CMs were fixed in 4% paraformaldehyde and permeabilized in 100% methanol. Cells were blocked in 5% normal goat serum ^5^ and incubated with primary antibodies diluted in 1% NGS: anti-NKX2-5 (Cell Signaling Technologies, #8792, 1:500), anti-cardiac troponin T (cTnT; Thermo Fisher Scientific, MA5-12960, 1:300), and anti-ATP5A (Abcam, ab176569, 1:200). Cells were then incubated with species-appropriate secondary antibodies (goat anti-mouse 488 (Southern Biotech, 103130, 1:200) and donkey goat anti-rabbit 594 (Sigma, SAB4600107, 1:200)) diluted in 1% NGS. Slides were mounted with ProLong Gold Antifade Mountant with DAPI (Thermo Fisher Scientific, P36931) and imaged using a Leica Stellaris 8 confocal microscope.

For analysis of sarcomere organization, hiPSC-derived cardiomyocytes stained for cTnT and NKX2-5 were classified into three categories based on myofibril organization: class 1, poorly organized myofibrils; class 2, intermediate organization; and class 3, highly organized, striated myofibrils. For analysis of mitochondrial distribution, hiPSC-derived cardiomyocytes stained for ATP5A were classified into three categories based on intracellular mitochondrial localization: class 1, predominantly perinuclear; class 2, intermediate; and class 3, predominantly cytoplasmic. For both analyses, cardiomyocytes were identified based on cTnT and/or NKX2-5 positivity, and only single, non-overlapping cells were included for quantification. All scoring was performed in a blinded manner.

### RNA-sequencing and analysis

For RNA-seq of embryonic hearts, total RNA was extracted using the miRNeasy Mini Kit (Qiagen, 217084) and quantified by NanoDrop spectrophotometry (Thermo Fisher Scientific). RNA integrity was assessed using a 2100 Bioanalyzer (Agilent Technologies) at the Emory Integrated Genomics Core. Poly(A)-selected RNA-seq libraries were prepared using the NEBNext Ultra II RNA Library Prep Kit (New England Biolabs), and paired-end sequencing (100 bp) was performed at 50 million reads per sample by Discovery Life Sciences (Huntsville, AL).

For RNA-seq of hiPSC-CMs, total RNA was extracted using the Aurum™ Total RNA Mini Kit (BioRad, 7326820). RNA quantification and integrity assessment were performed at the Emory Integrated Genomics Core using NanoDrop spectrophotometry (Thermo Fisher Scientific) and a 2100 Bioanalyzer (Agilent Technologies), respectively. Poly(A)-selected RNA-seq library preparation and paired-end sequencing (150 bp; 50 million reads per sample) were performed by Novogene (Sacramento, CA).

RNA-seq analysis was conducted using Partek Flow software (v11.0; Illumina). Raw sequencing reads were filtered and trimmed to remove low-quality bases and sequencing artifacts using Partek Flow quality-based trimming parameters, which scan reads from the 5′ and/or 3′ ends and trim bases below a defined quality threshold. Filtered reads were aligned to the mouse (GRCm39/mm39) or human (GRCh38.p14/hg38) reference genomes using STAR (v2.7.8a) ^66^. Read counts were generated using the Partek expectation–maximization (E/M) algorithm and normalized for differential gene expression analysis using the Partek Flow native implementation of DESeq2 ^67^. Unsupervised hierarchical clustering and heat map visualization were performed in Partek Flow. Differentially expressed genes were subjected to gene ontology and pathway enrichment analyses using Partek Flow, and gene ontology terms from embryonic heart datasets were summarized and visualized using Revigo ^25^. Differentially expressed mRNAs from E17.5 hearts and day 15 hiPSC-CMs were analyzed with the Metascape engine to determine ontology enrichment overlap between datasets ^44^.

### Flow Cytometry

hiPSC-derived cardiomyocytes were harvested, stained with ethidium monoazide (EMA), fixed in 4% paraformaldehyde, and resuspended in staining buffer. Cells were permeabilized in 90% methanol and blocked in blocking buffer. Cardiomyocytes were incubated with primary antibodies diluted in blocking buffer: anti-NKX2-5 (Cell Signaling Technologies, #8792, 1:800), anti-cardiac troponin T (Thermo Fisher Scientific, MA5-12960, 1:300), and anti-ATP5A (Abcam, ab176569, 1:100), followed by incubation with species-appropriate secondary antibodies (goat anti-mouse 488 (Southern Biotech, 103130, 1:200)), and goat anti-rabbit 594 (Sigma, SAB4600107, 1:200)). Fluorescent labeling was analyzed using a FACSymphony A5 flow cytometer (BD Biosciences).

### Immunoblotting

Total protein extracts were prepared from embryonic hearts, undifferentiated hiPSCs, and hiPSC-derived cardiomyocytes by solubilization in radioimmunoprecipitation assay (RIPA) buffer supplemented with a combined protease and phosphatase inhibitor cocktail (Thermo Fisher Scientific). Proteins were separated on 10% SDS–PAGE gels (TGX stain-free gels, Bio-Rad), transferred to PVDF membranes, and immunoblotted with the indicated antibodies. Membranes were imaged using a ChemiDoc XRS+ system (Bio-Rad).

Primary antibodies used were: anti-SLC25A1 (Proteintech, 15235-1-AP, 1:1000), OXPHOS Blue Native WB Antibody Cocktail (Abcam, ab110412, 1:500), anti-Alpha Tubulin (Cell Signaling Technologies, #2144, 1:1000) and anti-GAPDH (Fitzgerald, 10 R-G109A, 1:10,000). Secondary antibodies used were horseradish peroxidase-linked goat anti-mouse IgG (Cell Signaling Technologies, 7076), and horseradish peroxidase-linked goat anti-rabbit IgG (Cell Signaling Technologies, 7074).

### Mitochondrial respiration assays

Mitochondrial respiration in embryonic hearts was measured using an Oxygen Consumption Rate Assay Kit (Cayman Chemical, 600800) according to the manufacturer’s instructions.

Briefly, E17.5 hearts were isolated and minced in assay medium (DMEM lacking glucose supplemented with sodium pyruvate), a phosphorescent probe provided with the kit was added, and oxygen consumption rates (OCR) were measured using a fluorescence-based detection method. Changes in probe fluorescence over time were used to calculate extracellular OCR using dual-read ratiometric lifetime measurements on a Synergy Neo2 plate reader (BioTek). OCR values were normalized to protein content.

Mitochondrial respiration in hiPSC-derived cardiomyocytes was measured using a Seahorse XF Pro Analyzer (Agilent). Cardiomyocytes were seeded onto Matrigel-coated Seahorse XF96 plates and cultured for 72 h prior to analysis. On the day of the assay, cells were incubated in XF-RPMI medium supplemented with glucose, glutamine, and sodium pyruvate and equilibrated in a non-CO₂ incubator. Basal, proton leak, maximal, and non-mitochondrial respiration rates were determined by sequential injection of 1 μM oligomycin, 0.25 μM FCCP, and 0.5 μM rotenone/antimycin A. OCR were normalized to protein content.

### Statistics

The number of replicates for each experiment, as well as the numbers of mice/embryos used are denoted in figure legends. Unless stated otherwise, data are expressed as mean ± standard error of the mean. One-tailed Student’s t-test was used for comparisons of two groups and one-way ANOVA followed by post-hoc Dunnett’s test was used for multiple group comparisons tests. P < 0.05 were considered statistically significant and the following statistical significance indicators are used: ns denotes not significant, *P < 0.05; **P<0.01; \*\*\**P* < 0.001; ****P < 0.0001.

### Study approval

All animal experiments were conducted under protocols approved by Emory University’s Institutional Animal Care and Use Committee and in compliance with ethical regulations for animal use.

## RESULTS

### Partial loss of SLC25A1 does not impair placental development

Given the critical role of placental trophoblasts in mediating nutrient exchange and storage, as well as supporting embryonic cardiac development ^16,22–24^, we hypothesized that partial loss of SLC25A1 (*Slc25a1^+/-^*) may produce placental abnormalities similar to those reported in SLC25A1 homozygous deletion (*Slc25a1^-/-^*) embryos. Using our systemic *Slc25a1* knockout (KO) mouse line, we performed timed matings of *Slc25a1^+/-^* mice and collected placentas at embryonic day (E) 17.5 ^14^. Histological analyses of hematoxylin and eosin-stained placental sections were used to delineate the decidua, junctional zone, and labyrinth layers (Figure 1A). Consistent with previous reports, we found that *Slc25a1^-/-^* placentas exhibited a significant reduction in the junctional zone–to–total placental thicknesses as compared to *Slc25a1^+/+^*controls (Figure 1B). Notably, we also observed that *Slc25a1^-/-^* placentas displayed increased labyrinth-to-total thickness (Figure 1C), suggesting compensatory remodeling of placental architecture. Importantly, despite the presence of CHDs in *Slc25a1^+/-^* embryos as we previously reported ^14^, both junctional zone and labyrinth layer thicknesses were unchanged in *Slc25a1^+/-^*placentas relative to controls (Figure 1B,C), indicating that partial loss of SLC25A1 does not alter placental morphology or development.

**Figure 1.**
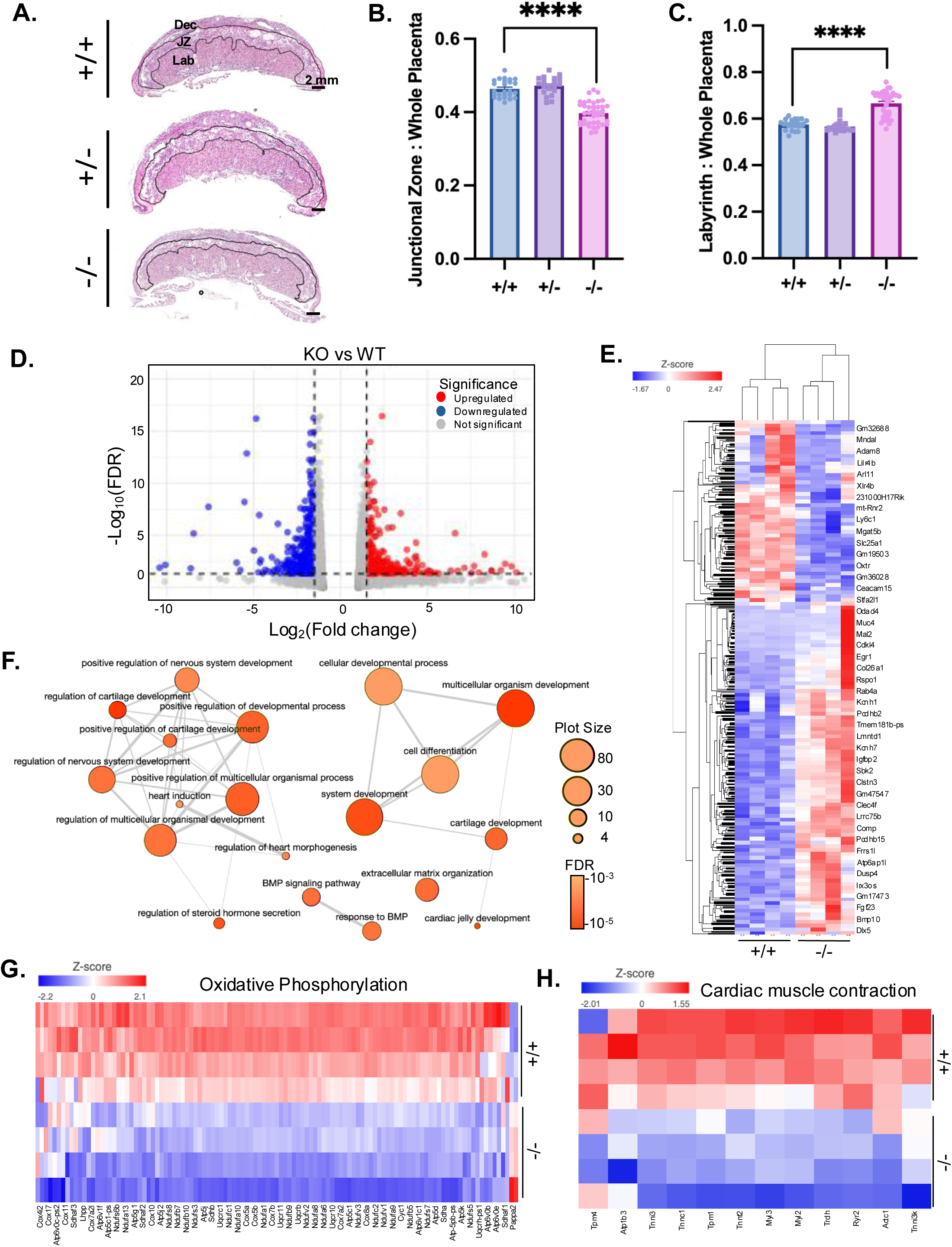
Systemic *Slc25a1* deletion dysregulates genes critical for cardiac morphogenesis and cardiac cell differentiation. A) Representative images of placental cross sections from *Slc25a1^+/+^*, *Slc25a1^+/-^*, *Slc25a1^-/-^* E17.5 embryos stained with hematoxylin and eosin. Dec indicates decidua, JZ indicate junctional zone, and Lab is the labyrinth. Junctional zone is outlined in black. Quantification of B) junctional zone to whole placenta ratios and C) labyrinth to whole placenta ratios in E17.5 placenta (n=22-40/group). D) Volcano plots comparing gene expression of *Slc25a1^-/-^* and *Slc25a1^+/+^* E17.5 hearts. Blue denotes downregulated transcripts and red denotes upregulated transcripts. E) Hierarchical clustering depicting DEGs in *Slc25a1^+/+^* vs. *Slc25a1^-/-^* E17.5 hearts. F) Gene ontologies identified with Revigo from DEGs in *Slc25a1^-/-^* versus *Slc25a1^+/+^* hearts^25^. Bubble color indicates the user-provided FDR; bubble size indicates the plot size. G) Heatmaps visualizing gene expression of oxidative phosphorylation and cardiac muscle contraction genes in E17.5 *Slc25a1^+/+^* vs. *Slc25a1^-/-^* hearts. Values reported as mean ± SEM. Kolmogorov-Smirnov test was used for statistical analysis. \*\*\*\**P* < 0.0001.

### Systemic *Slc25a1* deletion dysregulates genes critical for cardiac morphogenesis and cardiac cell differentiation

*Slc25a1^+/-^* embryos lacked placental developmental abnormalities (Figure 1B,C), but display CHDs ^14^, suggesting that placental derangements do not fully contribute to SLC25A1-dependent cardiac defects. Given that SLC25A1 is highly expressed in the developing myocardium and functions as a key regulator of metabolic maturation in the embryonic heart ^14^, we performed bulk RNA sequencing on E17.5 *Slc25a1^-/-^* and *Slc25a1^+/+^* embryonic hearts to identify transcriptional pathways dysregulated with SLC25A1 deletion.

Transcriptome analysis revealed 353 differentially expressed genes (DEGs; fold change ≥ 1.5, *Padj* < 0.05), 224 upregulated and 129 downregulated, between *Slc25a1^-/-^* and *Slc25a1^+/+^* hearts (Figure 1D). Hierarchical clustering of DEGs demonstrated distinct expression patterns between genotypes (Figure 1E). Gene ontology (GO) analysis of DEGs summarized using Revigo revealed enrichment of biological processes involved in heart induction, cardiac morphogenesis, and regulation of cardiac cell differentiation (Figure 1F)^25^.

Given the significant dysregulation of cardiac induction and morphogenetic pathways, we next examined the effect of SLC25A1 deletion on cardiac maturation gene programs. Heatmap visualization of gene from oxidative phosphorylation and cardiac muscle contraction gene ontologies demonstrated reduced expression of pathway-associated genes in *Slc25a1^-/-^* hearts relative to *Slc25a1^+/+^* controls (Figure 1G,H). Both pathways are critical markers of cardiac metabolic and structural maturation^26–29^. Together, these data demonstrate that systemic loss of SLC25A1 disrupts transcriptional programs governing cardiac morphogenesis and maturation, supporting a critical role for SLC25A1 in the regulation of developmental and metabolic gene networks in the heart.

### Cardiomyocyte-specific deletion of SLC25A1 in the developing heart produces cardiac structural and mitochondrial defects

To assess the cardiomyocyte-specific requirements for SLC25A1 in the developing heart, we generated mice with the conditional deletion of *Slc25a1* by crossing *Slc25a1* floxed mice (*Slc25a^fl^*^/*fl*^) with *Tnnt2-Cre* mice (which express Cre under the control of the cardiac troponin T promoter), which initiate recombination in cardiomyocytes beginning at E7.5 (*Slc25a1^fl/fl^;Tnnt2^Cre/+^*) (Figure 2A). Immunohistochemistry confirmed loss of SLC25A1 protein in the myocardium of *Slc25a1^fl/fl^;Tnnt2^Cre/+^* hearts at E14.5 (Figure 2B). At this developmental stage, when septation and ventricular wall compaction are largely complete ^3,4^, hearts from *Slc25a1^fl/fl^;Tnnt2^Cre/+^* embryos displayed significantly increased traceular myocardial thickness relative to *Slc25a1^fl/fl^* controls (Figure 2C,D), demonstrating that SLC25A1 deletion in developing cardiomyocytes causes cell-autonomous ventricular noncompaction.

**Figure 2.**
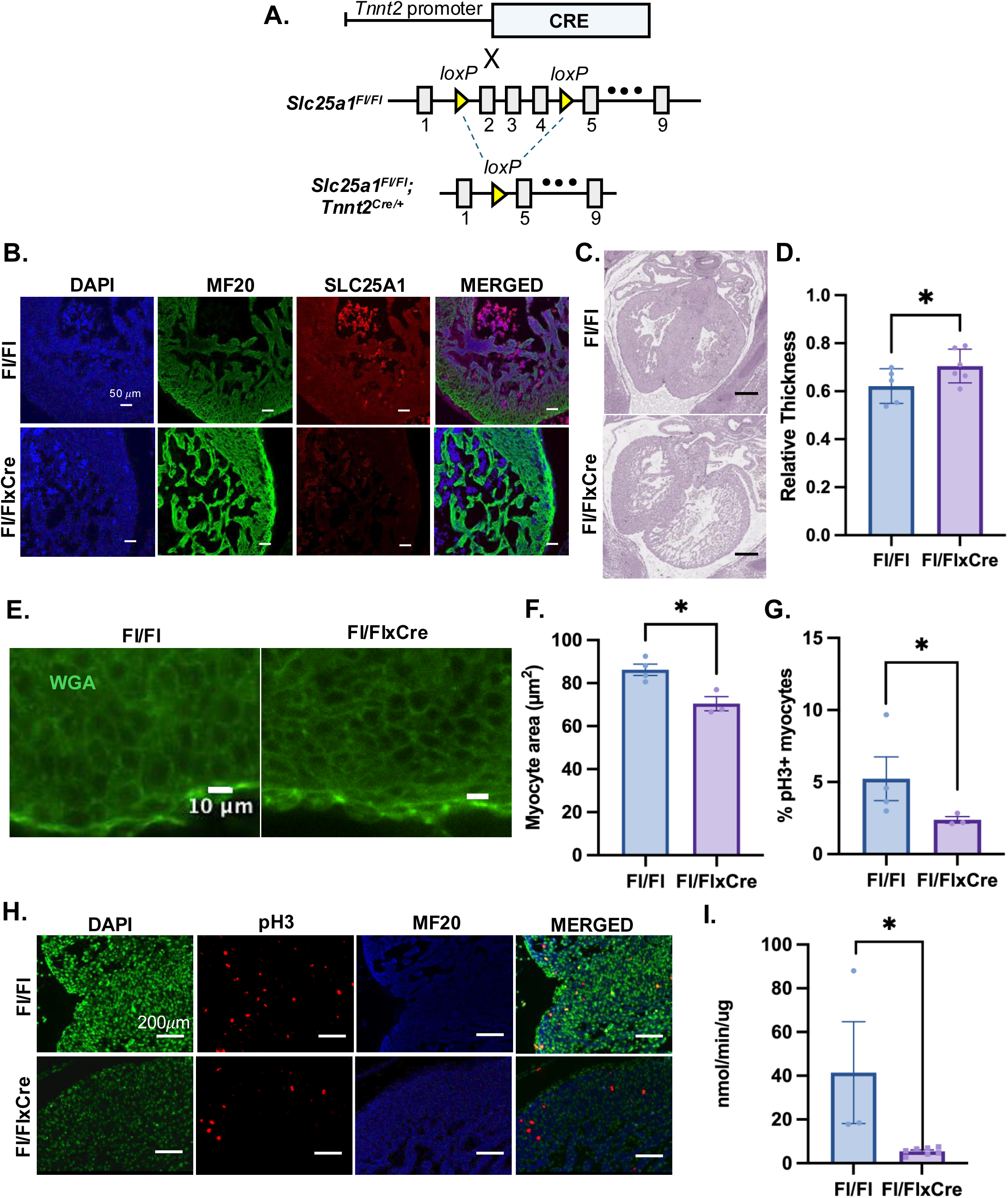
Cardiomyocyte-specific deletion of SLC25A1 in the developing heart produces cardiac structural and mitochondrial defects. A) Generation of *Slc25a1^fl/fl^;Tnnt2^Cre/+^* mice. B) Representative immunofluorescence images of SLC25A1 expression in E14.5 hearts from *Slc25a1^fl/fl^* and *Slc25a1^fl/fl^;Tnnt2^Cre/+^* embryos. DAPI was used to counterstain nuclei, and MF20 was used to counterstain cardiomyocytes. C) Representative images of E14.5 embryos from the indicated genotypes stained with hematoxylin and eosin. D) Quantification of relative trabecular myocardium thickness E14.5 hearts (n = 5/group). E) Representative immunofluorescence images of myocyte cell membranes counterstained with wheat germ agglutinin (WGA) in E14.5 hearts from the indicated genotypes. F) Quantification of myocyte area in E14.5 hearts (n = 3/group). G) Quantification of myocyte proliferation in E14.5 hearts (n = 3-4/group). H) Representative immunofluorescence images of pH3 expression in E14.5 heart from the indicated genotypes. DAPI was used to counterstain nuclei, pH3 was used to counterstain proliferative cells, and MF20 was used to counterstain cardiomyocytes. I) Quantification of mitochondrial oxygen consumption rates in E14.5 hearts (n = 3-7/group). Values reported as mean ± SEM. Student’s t-test was used for statistical analysis. *P < 0.05.

Proper ventricular maturation depends on a balance between cardiomyocyte growth, proliferation, and survival ^30–32^.To determine whether the morphologic defects we observed in *Slc25a1^fl/fl^;Tnnt2^Cre/+^* hearts were due to disruptions in these processes, we examined E14.5 hearts for cardiomyocyte size, proliferation, and apoptosis. Wheat germ agglutinin staining of the plasma membrane revealed decreased cardiomyocyte cross-sectional area in *Slc25a1^fl/fl^;Tnnt2^Cre/+^* hearts as compared to *Slc25a1^fl/fl^* controls (Figure 2E,F). Quantification of cardiomyocyte proliferation, assessed by phospho-histone H3 (pHH3) staining, revealed pHH3 positive nuclei in *Slc25a1^fl/fl^;Tnnt2^Cre/+^* hearts were significantly reduced (by 54%) as compared to controls, while TUNEL analysis showed no change in cell death with SLC25A1 deletion (Figure 2G,H; Figure S1). Taken together, SLC25A1 loss in cardiomyocytes causes decreased cardiomyocyte growth and proliferation in the developing heart.

Our previous work showed that systemic loss of SLC25A1 impairs mitochondrial maturation and respiration in the embryonic heart ^14^. Consistent with these findings, our transcriptomic analysis of Slc25a1^-/-^ hearts revealed downregulation of OXPHOS genes (Figure 1G). To determine whether mitochondrial dysfunction also occurs in the absence of SLC25A1 specifically within cardiomyocytes, we measured mitochondrial respiration in hearts from E17.5 *Slc25a1^fl/fl^;Tnnt2^Cre/+^* and *Slc25a1^fl/fl^* control embryos. *Slc25a1^fl/fl^;Tnnt2^Cre/+^* hearts displayed significantly reduced mitochondrial oxygen consumption rates as a compared to controls (Figure 2I), indicating an intrinsic requirement for SLC25A1 in cardiomyocytes for proper mitochondrial function.

### SLC25A1 is required cell-autonomously for cardiomyocyte differentiation

To further investigate the cell-autonomous roles of SLC25A1 in cardiomyocyte differentiation, we generated SLC25A1 KO human induced pluripotent stem cell (hiPSC) lines by CRISPR/Cas9-mediated genome editing using the wild type (WT) A18945 hiPSC line derived from CD34+ cord blood (Figure S2A) ^33^. Undifferentiated WT and SLC25A1 KO hiPSCs expressed pluripotency markers SSEA4, SOX2, TRA-1-81, and OCT4 (Figure S2B) and exhibited normal karyotypes (Figure S2C) ^34,35^. Trilineage differentiation confirmed their ability to form ectoderm (Nestin, PAX6) ^36,37^, mesoderm (NCAM, Brachyury) ^38,39^, and endoderm (CXCR4, SOX17)^40,41^ lineages (Figure S2D–F). Immunoblotting verified loss of SLC25A1 in two independent KO clones (KO1 and KO2; Figure 3A).

**Figure 3.**
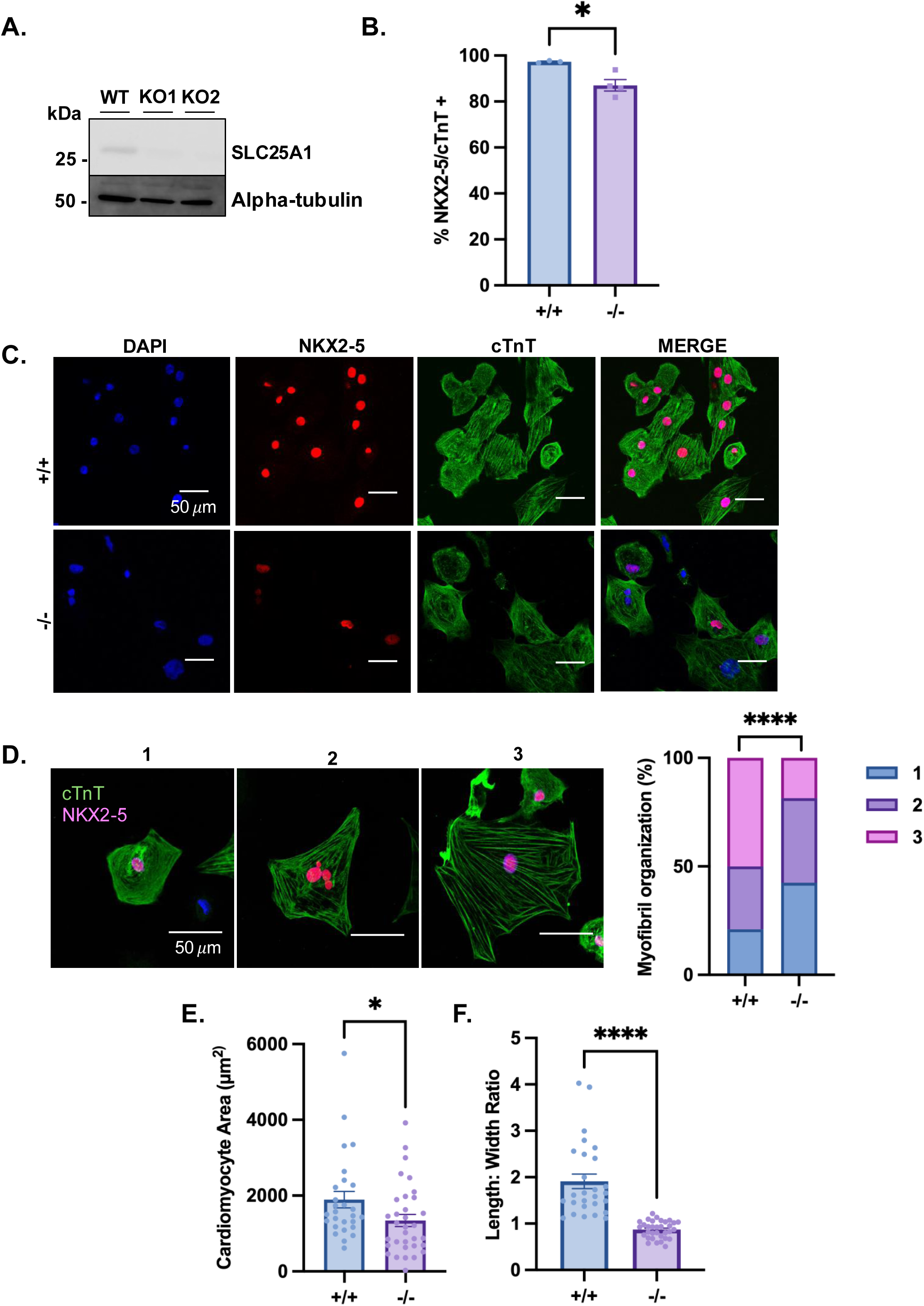
SLC25A1 is required cell-autonomously for cardiomyocyte differentiation. A) Western blot of SLC25A1 expression in total protein isolated from undifferentiated hiPSC cell lines. Alpha-tubulin staining was a protein loading control. B) Quantification of double positive NKX2-5 and cTnT rates from D15 hiPSC-CMs of indicated genotypes (n= 3-4 differentiations/group). C) Representative immunofluorescence images of NKX2-5 and cTnT expression in day 15 hiPSC-CMs from the indicated genotypes. DAPI was used to counterstain nuclei. D) Representative immunofluorescence of cTnT ^68^ and NKX2-5 (pink) staining in day 15 WT hiPSC-CMs for myofibril organization scoring. 1 denotes poor myofibril organization, 2 denotes an intermediate organization, and 3 denotes highly organized myofibrils. Quantification of myofibril organization from indicated genotypes (n= 54-144 cardiomyocytes/group). Quantification of hiPSC cardiomyocyte E) area and F) length to width ratios from indicated genotypes (n= 27-33 cardiomyocytes/group). Values reported as mean ± SEM. Student’s t-test and Chi-squared test were used for statistical analysis. *P < 0.05, \*\*\*\**P* < 0.0001.

To assess effects of SLC25A1 loss on cardiac differentiation, hiPSCs were differentiated into cardiomyocytes (hiPSC-CMs) by small molecule guided differentiation and analyzed on day (D) 15 of differentiation, a time point reflecting when cardiomyocyte identity is firmly established.

Flow cytometry analysis revealed reduced proportions of NKX2-5/cTnT double positive cells in SLC25A1 knockout lines as compared to WT controls (Figure 3B,C; Figure S3), suggesting a differentiation defect. Given similar differentiation phenotypes between clones, subsequent analyses focused on the KO1 line.

SLC25A1 KO hiPSC-CMs exhibited morphological abnormalities characterized by disorganized myofibrils and rounded cell shape. Myofibril organization was categorized into three classes (class 1, poorly organized; class 2, intermediate; and class 3, highly organized) and quantification revealed an increased proportion of cells with poor myofibril organization in KO hiPSC-CMs as compared to controls (Figure 3D). In addition, SLC25A1-KO hiPSC-CMs were smaller and rounder than controls, consistent with an immature cardiomyocyte morphology (Figure 3E,F) (25-27). Together, these findings suggest a cell-autonomous role for SLC25A1 in regulating cardiomyocyte differentiation.

### SLC25A1 deletion dysregulates gene programs required for cardiomyocyte differentiation and metabolic maturation

To define gene networks altered by SLC25A1 loss in differentiating cardiomyocytes, we performed bulk RNA sequencing on D15 WT and SLC25A1 KO hiPSC-CMs. From this analysis, 2766 DEGs (fold change ≥ 2, *Padj* < 0.05), 1337 upregulated and 1429 downregulated, were identified between KO and WT samples (Figure 4A). Hierarchical clustering revealed distinct transcriptional profiles between genotypes (Figure 4B). GO analysis of KEGG pathways revealed significant dysregulation of processes related to ventricular cardiac muscle morphogenesis (GO:0055010, FDR = 0.001), cardiocyte differentiation (GO:0035051, FDR =0.041), and cardiac muscle cell development (GO:0055013, FDR =0.049) (Figure 4C), consistent with the structural and differentiation defects observed in SLC25A1-deficient hearts and hiPSC-CMs (Figures 2 and 3). Expression of cardiomyocyte marker genes *TNNT2* (cardiac troponin T) and *MYH7* (beta-myosin heavy chain) were significantly reduced in KO hiPSC-CMs as compared to WT controls (Figure 4D,E) ^42^.

**Figure 4.**
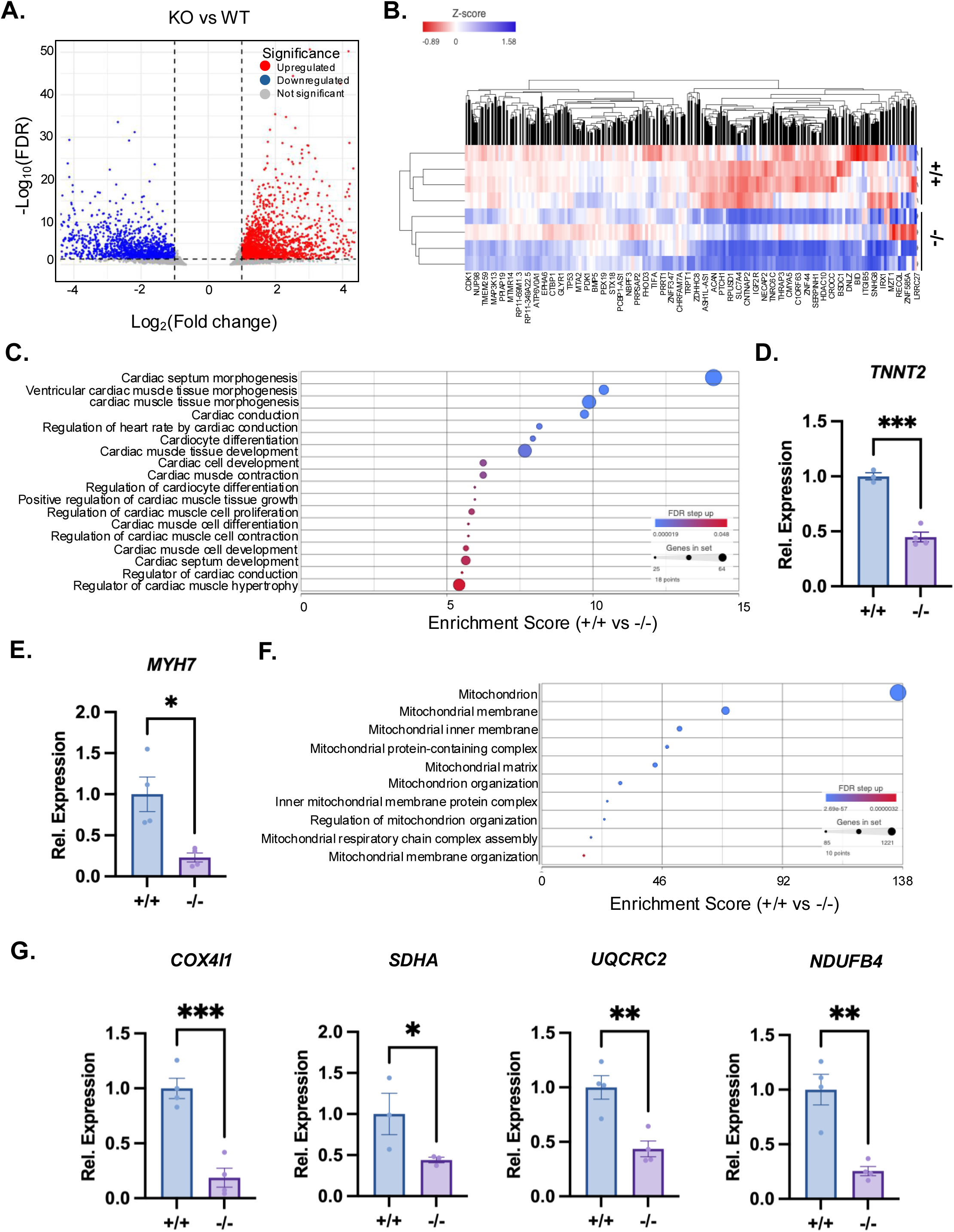
SLC25A1 deletion dysregulates gene programs required for cardiomyocyte differentiation and metabolic maturation. A) Volcano plot comparing gene expression of KO versus WT day 15 hiPSC-CMs. Blue denotes down-regulated transcripts and red denotes up-regulated transcripts. B) Heat map depicting differentially expressed genes in day 15 WT vs. KO hiPSC-CMs C) Top cardiac related gene ontologies identified with Partek Flow from genes differentially expressed in day 15 WT versus KO hiPSC-CMs. Quantification of relative transcripts expression levels of cardiomyocyte markers, D) *TNNT2* and E) *MYH7* gene on day 15 of hiPSC-CMs differentiation (n=4 differentiations/group). F) Top ten mitochondria related gene ontologies identified with Partek Flow from DEGs in WT versus KO hiPSC-CMs. Values reported as mean ± SEM. G) Quantification of relative expression levels for *COX4I1*, *SDHA*, *UQCRC2*, and *NDUFB4* in D15 WT and KO hiPSC-CMs (n=4 differentiations/group). Student’s t-test and weighted Kolmogorov-Smirnov test was used for statistical analysis. *P < 0.05; **P<0.01; \*\*\**P* < 0.001.

Because cardiomyocyte maturation depends on the establishing proper mitochondrial metabolic function, we next examined whether SLC25A1 loss affected gene programs related to mitochondrial organization and bioenergetics. Among the 45 mitochondria-related GO terms identified (Figure S4A), those associated with mitochondrial respiratory chain assembly and mitochondrial localization were significantly dysregulated in KO hiPSC-CMs (Figure 4F; Figure S4A). Additionally, expression of respiratory chain complex subunits *COX4I1*, *SDHA*, *UQCRC2*, and *NDUFB4* were significantly reduced in KO hiPSC-CMs as compared to WT controls (Figure 4D,E) ^43^.

Taken together, these results show that SLC25A1 loss disrupts transcriptional programs required for both cardiomyocyte differentiation and metabolic maturation.

### Conserved dysregulation of cardiomyocyte developmental gene programs across murine and human models upon SLC25A1 deletion

Using Metascape ^44^ to further analyze overlap between transcriptomic datasets from E17.5 mouse hearts and day 15 hiPSC-derived cardiomyocytes, we identified 62 differentially expressed genes (DEGs) that were commonly dysregulated upon SLC25A1 deletion (Figure 5A). Among these shared DEGs were *BMP10*, *BMP2*, *NPPA*, and *SLIT2*, all of which play established roles in cardiomyocyte differentiation and cardiac morphogenesis ^45–48^.

**Figure 5.**
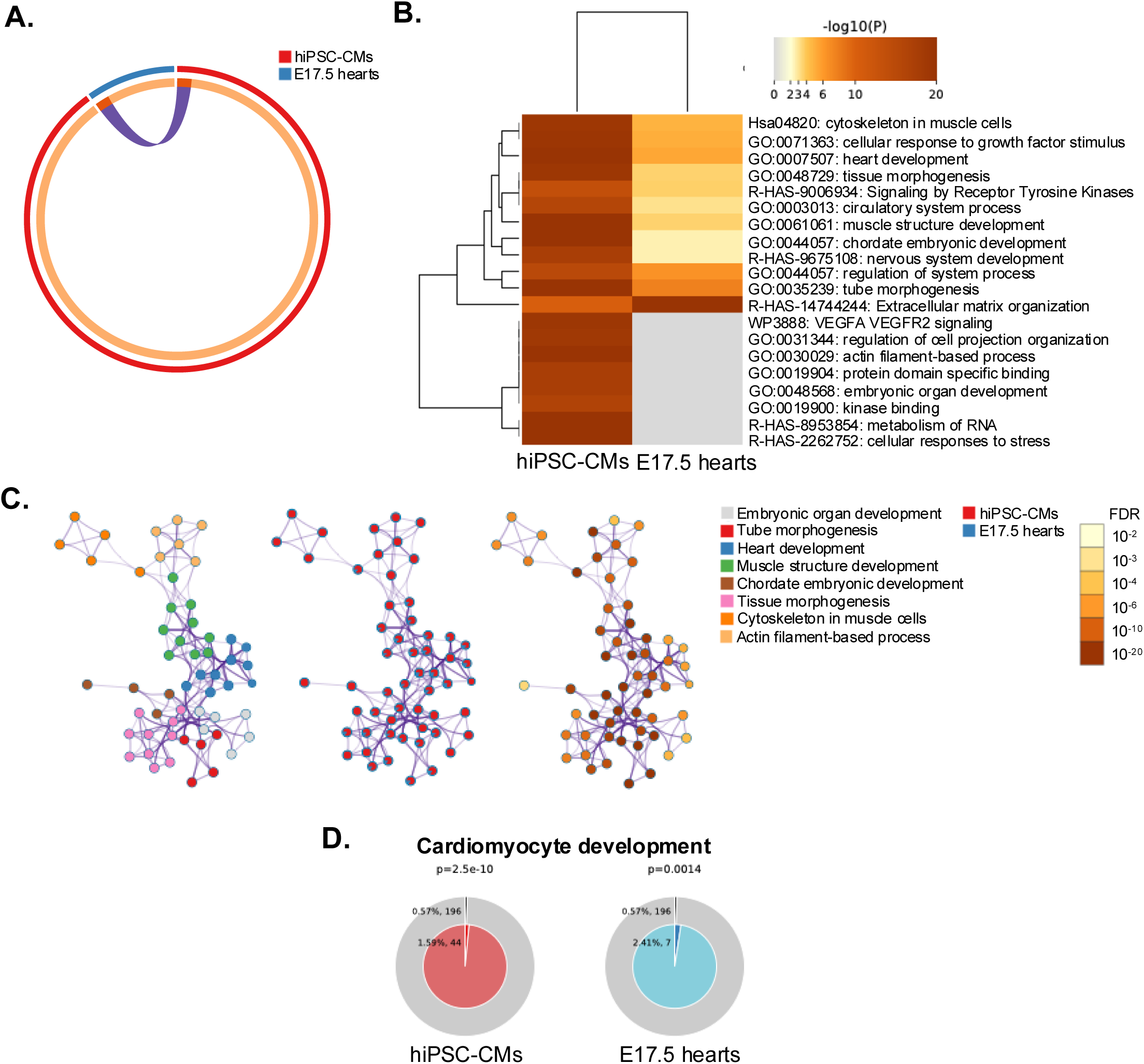
Conserved dysregulation of cardiomyocyte developmental gene programs across murine and human models upon SLC25A1 deletion. A) Overlap of differentially expressed genes (DEGs; n = 62) identified by RNA sequencing of day 15 hiPSC-derived cardiomyocytes and E17.5 embryonic mouse hearts following SLC25A1 deletion. B) Heatmap visualization of selected significantly enriched biological processes shared between day 15 hiPSC-derived cardiomyocytes and E17.5 embryonic hearts. C) Network visualization of enriched biological processes organized based on shared gene membership and enrichment significance across both models. D) Enrichment of cardiomyocyte development–associated genes among shared DEGs relative to genomic background in day 15 hiPSC-derived cardiomyocytes and E17.5 embryonic hearts.

To identify biological processes consistently perturbed across models, we performed pathway enrichment analysis and visualized significantly altered programs using a heatmap (Figure 5B). Prominent dysregulated pathways included cytoskeleton organization in muscle cells (hsa04820), heart development (GO:0007507), and extracellular matrix organization (R-HAS-14744244). These ontologies were further organized into Metascape-derived gene networks based on shared gene membership and enrichment significance (Figure 5C).

Given that cardiomyocyte development and differentiation were significantly disrupted upon loss of SLC25A1, we next examined whether gene programs underlying these processes were consistently dysregulated across models. Genes associated with cardiomyocyte development were significantly altered in both E17.5 hearts and day 15 hiPSC-derived cardiomyocytes following SLC25A1 deletion (Figure 5D).

Together, these analyses reveal a conserved set of cardiomyocyte developmental and morphogenetic gene programs that are disrupted by SLC25A1 loss across both embryonic mouse hearts and human hiPSC-derived cardiomyocytes.

### SLC25A1 deletion impairs mitochondrial function and organization in hiPSC-derived cardiomyocytes

We next asked whether the dysregulation of mitochondrial gene programs in SLC25A1-deficient hiPSC-CMs was accompanied by defects in respiratory chain complex expression. To assess mitochondrial respiratory chain integrity, we performed immunoblotting for core subunits of the respiratory complexes at day 20 (D20) of differentiation. SLC25A1 KO hiPSC-CMs exhibited reduced expression of complexes II-SDHA, IV-COXIV, and V-ATP5A subunits compared with WT controls (Figure 6A,B), consistent with transcriptomic evidence of disrupted OXPHOS gene expression in SLC25A1-deficient hiPSC-CMs and embryonic hearts.

**Figure 6.**
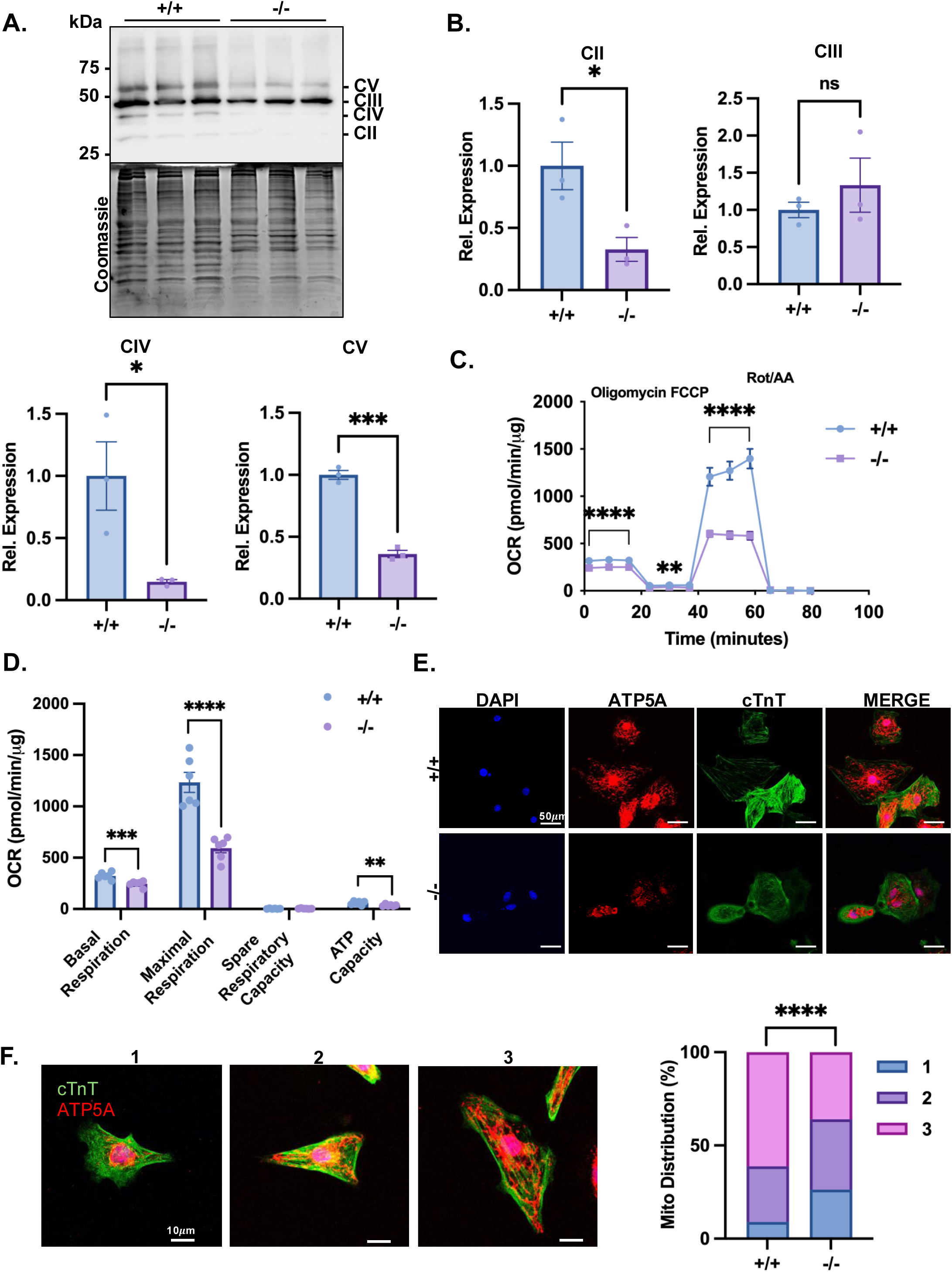
SLC25A1 deletion impairs mitochondrial function and organization in hiPSC-derived cardiomyocytes. A) Western blot of OXPHOS complex expression in total protein isolated from day 20 hiPSC-CM of indicated genotypes. Coomassie staining was used a protein loading control. B) Densitometry quantification of complex II, III, IV, and V expression (n = 3/group). C) Real time oxygen consumption rate (OCR) measurement of day 15 WT and KO iPSC-CMs sequentially treated with oligomycin, FCCP, and Rotenone/Antimycin A. D) Quantification of basal respiration, maximal respiration, spare respiratory capacity, ATP capacity (n = 6 differentiations/group). E) Representative immunofluorescence images of cTnT ^68^ and ATP5A (red) expression in day 15 hiPSC-CMs from the indicated genotypes. DAPI was used to counterstain nuclei. F) Representative immunofluorescence of cTnT ^68^ and ATP5A (red) staining demonstrating mitochondrial distribution in day 15 hiPSC-CMs for mitochondrial distribution scoring. 1 denotes perinuclear mitochondrial distribution, 2 denotes an intermediate mitochondrial distribution, and 3 denotes cytoplasmic mitochondrial distribution. Quantification of mitochondrial distribution from indicated genotypes (n= 67-114 cardiomyocytes /group). Values reported as mean ± SEM. Student’s t-test and Chi squared test were used for statistical analysis. ns denotes not significant; *P<0.05; **P<0.01; \*\*\**P* < 0.001; \*\*\*\**P* < 0.0001.

To determine the functional consequences of reduced respiratory complex expression, we measured mitochondrial oxygen consumption in SLC25A1-deficient and control hiPSC-CMs. In alignment with our findings in *Slc25a1^fl/fl^Tnnt2^Cre/+^* hearts and previously published data from systemic SLC25A1 deletion hearts ^14^, SLC25A1 KO hiPSC-CMs exhibited significantly lower basal and maximal respiration rates compared with WT cells (Figure 6C, D).

Because RNA-sequencing of SLC25A1-deficient hiPSC-CMs revealed dysregulation of mitochondrial gene programs, we next examined whether these transcriptional changes were reflected at the organelle level through alterations in mitochondrial organization and function. Because GO analysis revealed dysregulation of mitochondrial localization pathways, we next examined mitochondrial distribution within hiPSC-CMs using immunostaining for the mitochondrial ATP synthase subunit ATP5A. SLC25A1 KO hiPSC-CMs displayed predominantly perinuclear mitochondrial localization, characteristic of immature cardiomyocytes, whereas WT cells exhibited more cytoplasmic mitochondrial networks (Figure 6E) ^49–52^. Quantification based on mitochondrial localization patterns (class 1, perinuclear; class 2, intermediate; class 3, cytoplasmic) showed a significant increase in class 1 cells and a decrease in class 3 cells upon SLC25A1 deletion (Figure 6F). Thus, SLC25A1 deletion alters mitochondrial organization toward an immature perinuclear pattern.

Collectively, these findings demonstrate that SLC25A1 is required for both mitochondrial distribution and respiratory function in hiPSC-CMs. Together with our in vivo data, these results establish that SLC25A1 functions cell-autonomously within cardiomyocytes to support mitochondrial and cardiac maturation.

## DISCUSSION

Cardiac morphogenesis requires the coordination of mitochondrial and metabolic maturation with cardiomyocyte differentiation. Yet, how these processes are mechanistically linked during development remains incompletely understood. In this study, we identify a previously unrecognized cell-autonomous role for SLC25A1 in cardiomyocyte differentiation and mitochondrial maturation. By integrating systemic and cardiomyocyte-specific SLC25A1 KO mice with SLC25A1 KO hiPSC-CMs, we show that SLC25A1 functions within cardiomyocytes to support mitochondrial organization and function, bioenergetic maturation, and proper ventricular development, and defines a metabolic axis by which mitochondrial function controls cardiac morphogenesis.

The placenta-fetal heart axis is a well-established regulator of cardiac development, and prior studies have implicated SLC25A1 in placental morphogenesis and placental-driven cardiac defects ^16,18,24^. Consistent with these reports, we observed placental abnormalities in *Slc25a1^-/-^*embryos. However, hemizygous loss of *Slc25a1* produced CHDs in the absence of overt placental malformations, indicating that placental dysfunction is not required for SLC25A1 deletion-associated cardiac phenotypes ^14^. Notably, we also observed compensatory remodeling of placental architecture in *Slc25a1^-/-^* placentas, where the junctional zone reduction was accompanied by an expansion of the labyrinth layer ^53^. While this suggests that the placenta can remodel to adapt to SLC25A1 loss, our data also suggests that these changes are not sufficient to prevent SLC25A1 deletion-associated CHD, indicating that SLC25A1 is needed beyond the placenta for proper cardiac morphogenesis.

Although previous studies using cardiomyocyte-specific *Slc25a1* deletion models reported no overt postnatal cardiac abnormalities ^16^, embryonic phenotypes were not examined in detail. Here, we show that deletion of *Slc25a1* in developing cardiomyocytes disrupts ventricular wall compaction, cardiomyocyte growth and proliferation, and mitochondrial respiration-phenotypes that parallel those observed in systemic KO hearts. These results establish a cell-autonomous requirement for SLC25A1 in cardiomyocyte maturation during heart development. Notably, cardiomyocyte-specific deletion does not fully recapitulate the spectrum of CHDs observed in systemic KOs, suggesting that SLC25A1 may also function in additional cardiac or extra-cardiac cell types to coordinate overall heart morphogenesis.

Our hiPSC-CM studies provide direct support for a cardiomyocyte-intrinsic role of SLC25A1. SLC25A1-KO hiPSCs exhibit impaired differentiation, immature cellular morphology, disorganized myofibrils, and perinuclear mitochondrial localization which are hallmarks of immature cardiomyocytes ^49–52,54–58^. These defects were accompanied by dysregulation of genetic programs associated with OXPHOS, mitochondrial organization, and respiratory chain assembly, suggesting that SLC25A1-dependent mitochondrial citrate export regulates bioenergetic programs required for cardiomyocyte differentiation and maturation. Importantly, comparative transcriptomic analyses revealed that loss of SLC25A1 induces a conserved disruption of cardiomyocyte developmental and morphogenetic gene programs across both murine embryonic hearts and human hiPSC-derived cardiomyocytes, indicating that SLC25A1 regulates a core cardiomyocyte differentiation program that is preserved across species

Beyond insights into the mitochondrial control of heart development, these findings have implications for congenital heart disease in humans. SLC25A1 is compromised in 22q11.2 deletion syndrome (22q11DS) ^20,59,60^, a chromosomal microdeletion disorder that is one of the most common genetic causes of CHD^61^. Our prior work showed that *Slc25a1* deletion produces CHD resembling those observed in 22q11DS, and ultrarare pathogenic variants in *SLC25A1* are associated with CHDs that are also observed in 22q11DS ^14^. The present study refines this connection by identifying cardiomyocyte-intrinsic mitochondrial and metabolic dysfunction as a potential mechanism through which SLC25A1 contributes to cardiac malformations, and raises the possibility that *SLC25A1* may contribute in part to CHD observed in 22q11DS.

In summary, our findings position SLC25A1 as a cardiomyocyte-intrinsic regulator of mitochondrial maturation, metabolic remodeling, and cardiac differentiation. By linking mitochondrial metabolism to morphogenetic programs, SLC25A1 defines a metabolic axis that is essential for normal heart development.

## AUTHOR CONTRIBUTIONS

C.O. and J.Q.K. wrote the manuscript. C.O., N.A.A., N.G., E. H. L., N.T., M.J.H., A.H., and J.Q.K. performed experiments. J.Q.K., V.F., B.D., C.X., A.H., C.O., and N.A.A., analyzed the data. J.Q.K. and C.O. designed the study. J.Q.K. conducted experimental oversight.

## ACKNOWLEDGEMENTS

The authors thank Austin Park for assistance with animal husbandry and technical support. J.Q.K. was supported by the was supported by the Additional Ventures Single Ventricle Research Fund, the Saving Tiny Hearts Society, the Department of Defense (0000063651), and the National Institutes of Health (R01-GM-144729). C.O. was supported by an American Heart Association Predoctoral Fellowship (24PRE1199924). N.G. was supported by an American Heart Association Predoctoral Fellowship (25PRE1372965) and an ARCS Foundation Scholarship. C.X. was supported by the National Institutes of Health (R01AA028527 and R21CA285254). V.F. was supported by the National Institutes of Health (ES034796). This work was supported in part by the Emory University Stem Cell and Organoids Core (ESCOC; RRID:SCR_023264), the Emory Integrated Genomics Core (EIGC; RRID:SCR_023529), the Emory Flow Cytometry Core (EFCC; RRID:SCR_023536), and the Integrated Cellular Imaging Shared Resource of the Winship Cancer Institute of Emory University (NIH/NCI P30CA138292). These core facilities are supported in part by the Georgia Clinical and Translational Science Alliance of the National Institutes of Health under Award Number UL1TR002378.The content is solely the responsibility of the authors and does not necessarily represent the official views of the National Institutes of Health.

## FIGURE LEGENDS

**Supplemental Figure 1. Cardiomyocyte-specific deletion of *Slc25a1* does not alter myocyte cell death.** A) Representative immunofluorescence images of TUNEL positive cells in E14.5 heart from the indicated genotypes. DAPI was used to counterstain nuclei and TUNEL was used to counterstain dead cells. B) Quantification of myocyte cell death in E14.5 hearts (n = 3-4/group). Values reported as mean ± SEM. Student’s t-test was used for statistical analysis; ns denotes not significant.

**Supplemental Figure 2. hiPSC cell lines are undifferentiated and pluripotent.** A) CRISPR/Cas9 deletion strategy in generated hiPSC clones. B) Representative immunofluorescent images of expression of undifferentiation markers, SSEA4, SOX2, TRA-1-81, and Oct 4 in SLC25A1 hiPSC clones (F5-1 WT, B4-1 KO, and E2-1 KO). DAPI was used to counterstain nuclei. Scale bar is 20 µm. C) Representative Karyotype-G banding hiPSC clones F5-1 WT, B4-1 KO, and E2-1 KO hiPSC clones. D) Representative immunofluorescent images of expression of ectoderm markers, Nestin and Pax6 in ectoderm-differentiated hiPSC clones. Scale bar is 20 µm. E) Representative dot plots of flow cytometric analysis of expression of endoderm markers, CXCR4 and Sox17, in endoderm differentiated hiPSC clones. F) Representative dot plots of flow cytometric analysis of expression of mesoderm markers, NCAM and Brachyury, in hiPSC clones from mesoderm-differentiated cell lines.

**Supplemental Figure 3. SLC25A1 deletion in alternate cell line impairs hiPSC-CM differentiation.** A) Quantification of double positive NKX2-5 and cTnT rates from WT and KO2 D15 hiPSC-CMs (n= 3 differentiations/group). Values reported as mean ± SEM. Student’s t-test was used for statistical analysis. *P < 0.05.

**Supplemental Figure 4. S*L*C25A1 deletion dysregulates mitochondrial gene programs required in mature hiPSC-CMs.** A) Comprehensive gene ontologies analysis of mitochondria-related gene ontologies identified with Partek Flow from DEGs in WT versus KO hiPSC-CMs.

## ACKNOWLEDGEMENTS

The authors thank Austin Park for assistance with animal husbandry and technical support. J.Q.K. was supported by the was supported by the Additional Ventures Single Ventricle Research Fund, the Department of Defense (0000063651), and the National Institutes of Health (R01-GM-144729). C.O. was supported by an American Heart Association Predoctoral Fellowship (24PRE1199924). N.G. was supported by an American Heart Association Predoctoral Fellowship (25PRE1372965) and an ARCS Foundation Scholarship. C.X. was supported by the National Institutes of Health (R01AA028527 and R21CA285254). V.F. was supported by the National Institutes of Health (ES034796). This work was supported in part by the Emory University Stem Cell and Organoids Core (ESCOC; RRID:SCR_023264), the Emory Integrated Genomics Core (EIGC; RRID:SCR_023529), the Emory Flow Cytometry Core (EFCC; RRID:SCR_023536), and the Integrated Cellular Imaging Shared Resource of the Winship Cancer Institute of Emory University (NIH/NCI P30CA138292). These core facilities are supported in part by the Georgia Clinical and Translational Science Alliance of the National Institutes of Health under Award Number UL1TR002378.The content is solely the responsibility of the authors and does not necessarily represent the official views of the National Institutes of Health.

