## Supplementary figures and images for "Cardiomyocyte-intrinsic SLC25A1 regulates cardiac differentiation and mitochondrial function"

### Supplemental Figures

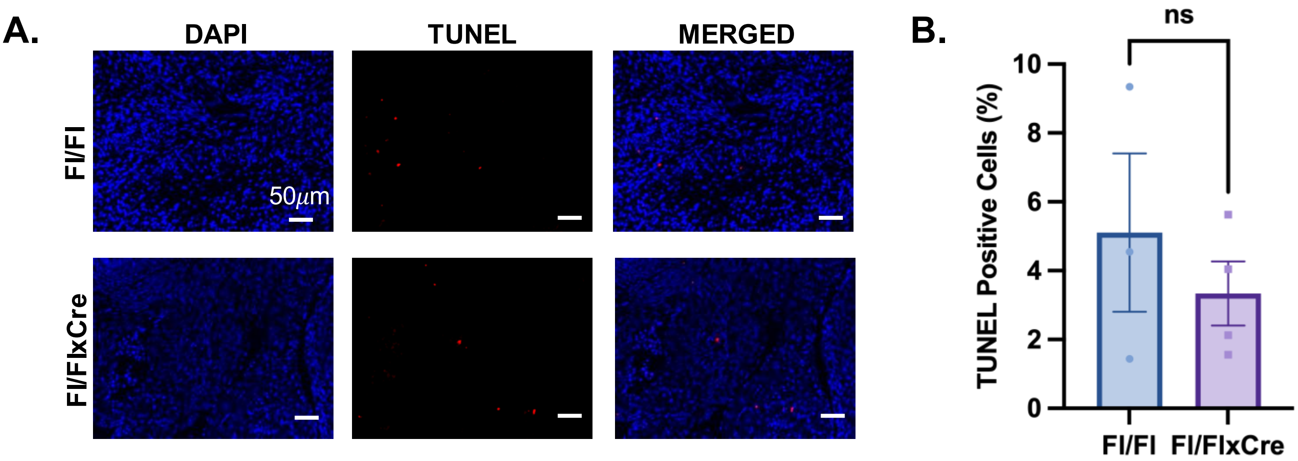

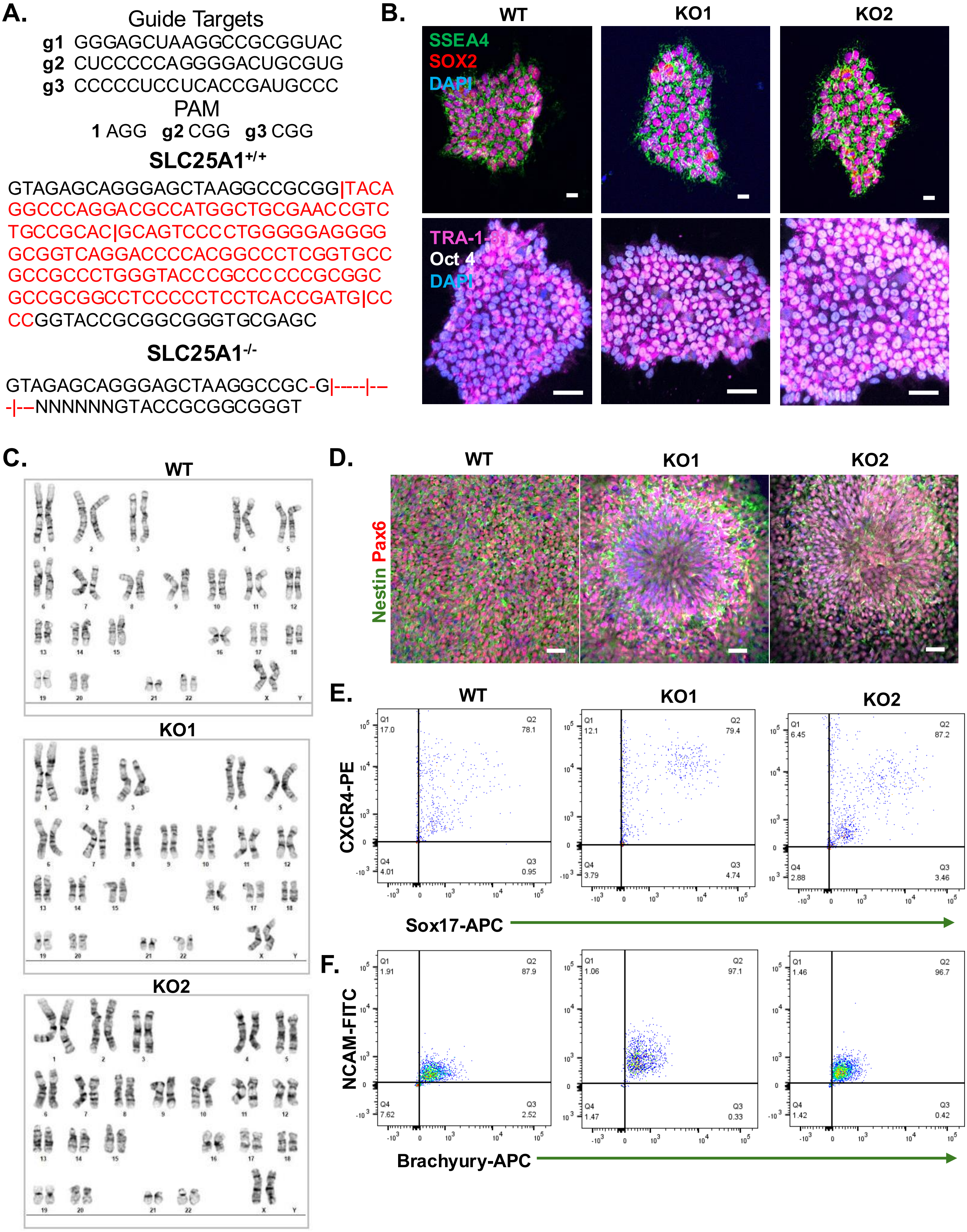

**Figure S3.**

**A.**

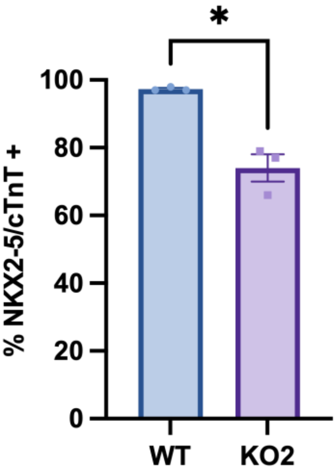

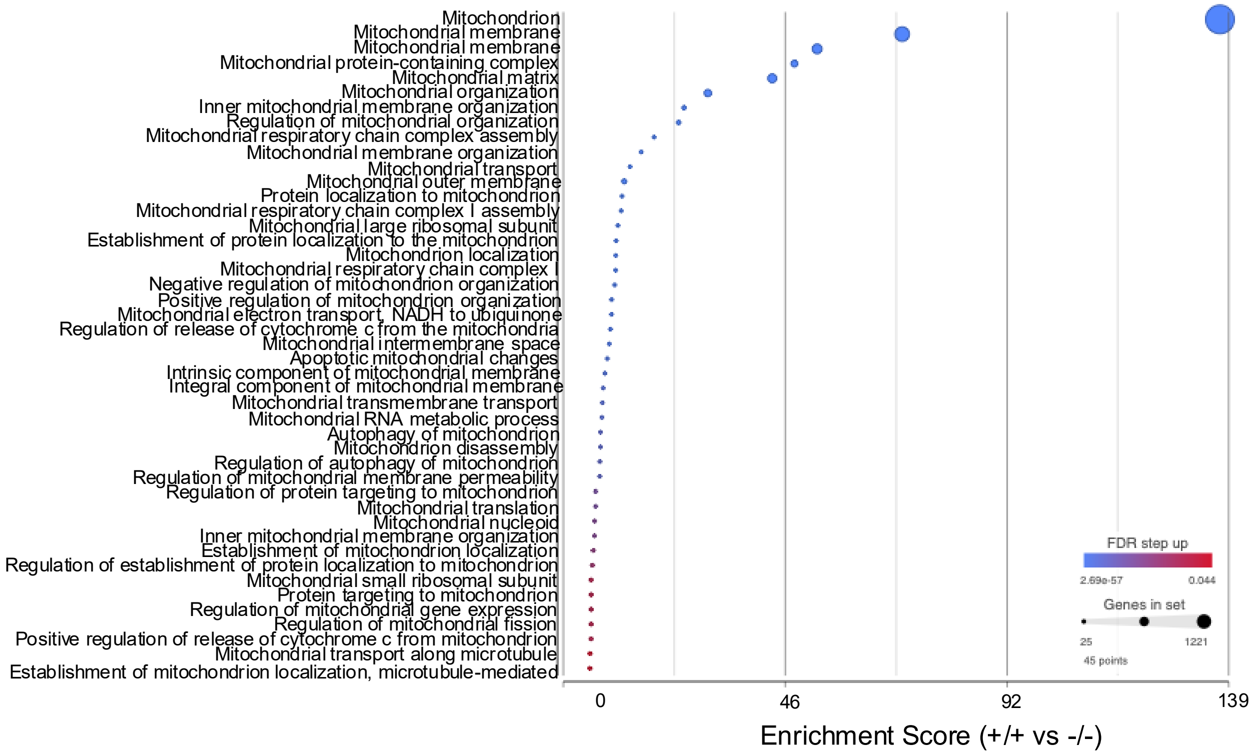
